# CytoGPS: A Web-Enabled Karyotype Analysis Tool for Cytogenetics

**DOI:** 10.1101/575423

**Authors:** Zachary B. Abrams, Lin Zhang, Lynne V. Abruzzo, Nyla A. Heerema, Suli Li, Tom Dillon, Ricky Rodriguez, Kevin R. Coombes, Philip R. O. Payne

**Affiliations:** Departments of Biomedical Informatics, Wexner Medical Center, The Ohio State University, Columbus, OH 43210 USA; Institute for Informatics, Washington University School of Medicine in St. Louis, St. Louis, MO 63108; Departments of Pathology, Wexner Medical Center, The Ohio State University, Columbus, OH 43210 USA

## Abstract

Karyotype data are the most common form of genetic data that is regularly used clinically. They are collected as part of the standard of care in many diseases, particularly in pediatric and cancer medicine contexts. Karyotypes are represented in a unique text-based format, with a syntax defined by the International System for human Cytogenetic Nomenclature (ISCN). While human-readable, ISCN is not intrinsically machine-readable. This limitation has prevented the full use of complex karyotype data in discovery science use cases. To enhance the utility and value of karyotype data, we developed a tool named CytoGPS. CytoGPS first parses ISCN karyotypes into a machine-readable format. It then converts the ISCN karyotype into a binary Loss-Gain-Fusion (LGF) model, which represents all cytogenetic abnormalities as combinations of loss, gain, or fusion events, in a format that is analyzable using modern computational methods. Such data is then made available for comprehensive “downstream” analyses that previously were not feasible.

**Availability and Implementation:** Freely available at https://cytogps.org

**Supplementary information:** Not applicable

## 1 Introduction

Karyotype data are the most common form of genetic data regularly employed in clinical medicine. They are stored in a text-based standard format with a syntax defined by the International System for human Cytogenetic Nomenclature or ISCN (McGowan-Jordan, et al., 2016). This international standard is updated regularly and dictates how karyotypes should be written and stored. Although trained cytogeneticists are able to read ISCN, the nomenclature is formatted as a semi-context-free and semi-structured language, thus making it incompatible with comprehensive computational analysis. This limitation explains why past attempts to parse karyotype data have only been marginally successful.

To make karyotype more readily available for computational analyses, we developed a rules-based grammar parser called CytoGPS. The parser identifies the relevant cytogenetic information in an ISCN-encoded karyotype, and then uses a mapping system to translate that information into a computer readable format, which we refer to as the Loss-Gain-Fusion (LGF) model (Abrams et al., 2015). In this note, we describe a website (http://cytogps.org) that we built for individual researchers so that they can quickly and easily use CytoGPS. The website provides both single karyotype and batch karyotype processing. It enables researchers to automatically process thousands of karyotypes written in the short form ISCN nomenclature. Further, CytoGPS supports the development of future, structured databases based on the LGF model, which we believe will further cytogenetic research by unlocking the potential currently locked within non-computable ISCN-encoded karyotypes.

## 2 Implementation

As was noted previously, the foundation of CytoGPS is a grammar-based parser that was created using Antlr, a system designed to construct grammar sets (Parr, 2013). Using this parser, CytoGPS can translate ISCN-encoded karyotypes into a parse tree that is capable of decomposing a given karyotype into subcomponents. The resulting parse tree can then be traversed to extract relevant information from different branches and leaves of the tree, thus extracting biological information from the ISCN karyotype.

When parsing a karyotype, the most important information that is encoded is the description of cytogenetic abnormalities located on specific chromosome bancs. This information can be thought of as the event and its location. These event/location pairs are translated by CytoGP using a rule-based mapper that contains an enumeration of all ISCN cytogenetic abnormalities as well as a biological interpretation of those abnormalities in the context of losses, gains, and fusions. This rules-based system ultimately translates the event/location information from the ISCN karyotype into a binary model that represents each such Loss-Gain-Fusion event (what we call the LGF model). In the LGF model, each cytogenetic band is represented three times – once each for loss, gain, and fusion, preserving the cytogenetic meaning of the karyotype.

For ease of use, we developed a website to allow researchers and clinicians the ability to access CytoGPS in an easy and highly usable manner (Fig. 1). To facilitate access to the tool, we made a few key design decisions. First, the homepage gives an overview of the purpose of CytoGPS its functionality. Users may then navigate to an interactive set of interface components, where more detailed descriptions about required inputs to the tools are provided. Third, we include detailed examples and example data sets in both the single karyotype analysis and batch karyotype analysis so that users can explore the system to determine whether CytoGPS can help them meet their research needs. All of these components provide clear visual feedback to the user on the options they have selected as well as how the single karyotype analysis results can be shown. Although we only provide data visualization for individual karyotype analysis, users can individually assess each affected chromosome within their input karyotype via an ideogram that presents each chromosome in horizontal view. By clicking on an individual chromosome, users can access a vertical view of that chromosome that displays more detailed information about each cytogenetic band (Fig. 1). This visually demonstrates how the CytoGPS algorithm translates the ISCN data into the LGF model. For batch karyotypes, we return a file containing the results for each individual karyotype in the batch along with statistics summarizing the entire cohort of karyotypes. These statistics include the percentage of the input population that had specific cytogenetic abnormalities, so users can easily determine the defining cytogenetic events for their data set.

**Figure 1:**
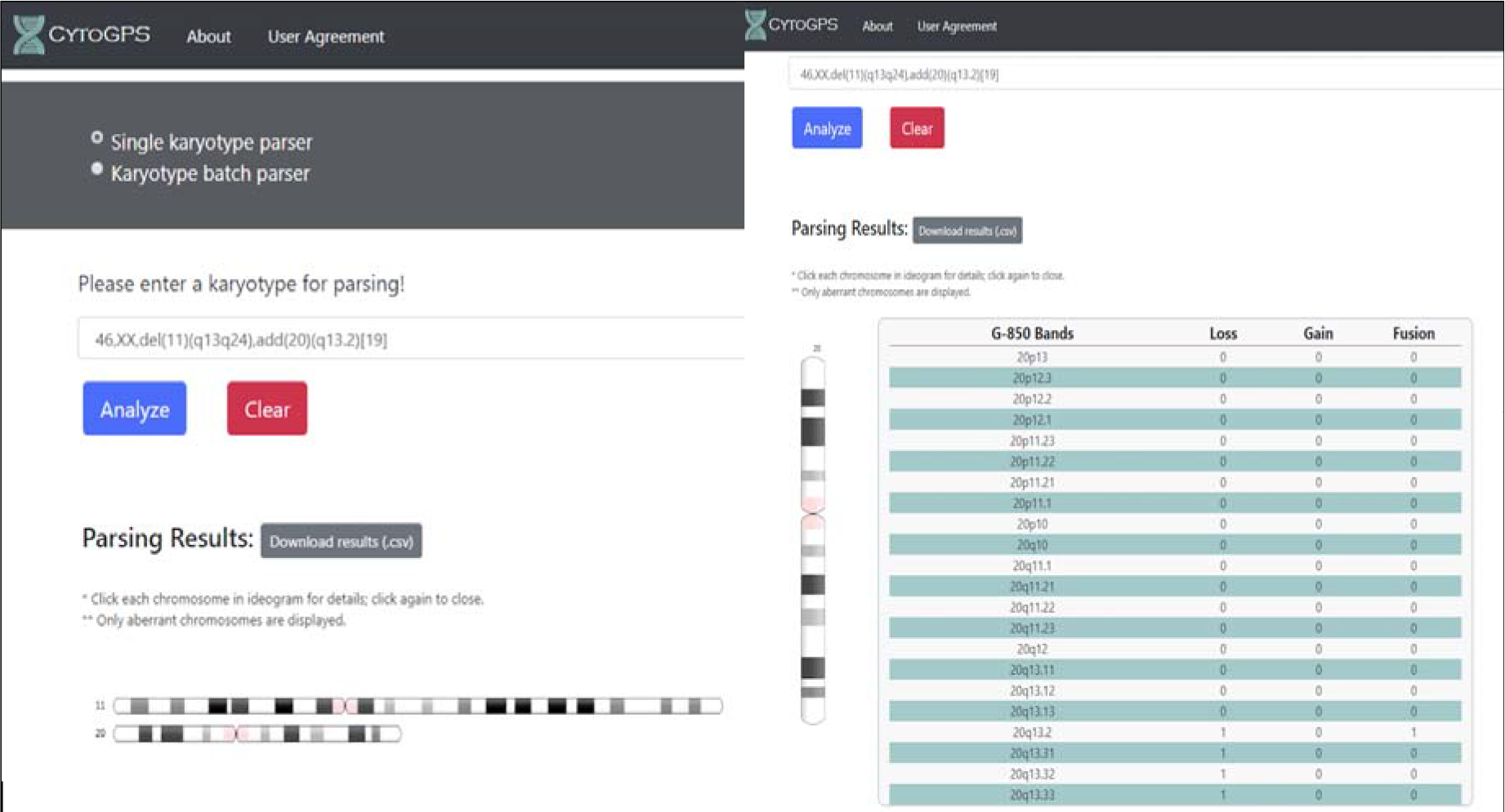
CytoGPS Website. This shows the karyotype analysis landing page, which describes general information about how to use the web based analysis tool. The left panel shows how results are displayed after a karyotype has been parsed. The user is shown which chromosomes have cytogenetic abnormalities and is provided with a downloadable file of the results. When a user clicks on a chromosome, they see a table in the right panel that shows each cytogenetic region on that particular chromosome and how those

## 3. Conclusion

CytoGPS is a unique research tool for the computational analysis of karyotype data. By developing a user-friendly and free website, we hope to improve the dissemination and usability of the CytoGPS algorithm to aid a wider number of research groups in analyzing their cytogenetic data.

## Funding

This work was supported by the National Library of Medicine (NLM) grant number T15 LM011270, the National Cancer Institute grant number R03 CA235101, and by Pelotonia Intramural Research Funds from the James Cancer Center.

